# Chronic infection of *Caenorhabditis elegans* by Orsay virus induces age-dependent immunity and superinfection exclusion

**DOI:** 10.1101/2024.12.26.630373

**Authors:** Victoria G. Castiglioni, Ana Villena-Giménez, Antonio González-Sánchez, Christina Toft, Gustavo G. Gómez, Santiago F. Elena

## Abstract

Orsay virus (OrV) is a natural pathogen of *C. elegans*, which mounts an antiviral response upon infection and enables the investigation of the mechanisms governing infection and immunity. Here, we focus on two of these features, namely the effect of life-long infections and superinfection dynamics. By following the course of an infection throughout the lifespan of a synchronous wild-type population, we describe several viral load peaks followed by sharp decreases in viral load, suggesting that the infection is chronic and that animals manage to suppress viral replication successfully throughout most of their lives. Moreover, we show that animals that have been previously exposed to the virus are able to control viral replication upon a subsequent inoculation, indicative of superinfection exclusion. Primary infections produced transcriptomic and small RNA alterations, whose extent was highly dependent on the developmental stage of the worm at the time of infection and the time until sampling. Superinfection, in turn, had little impact on the overall transcriptome, but showed a misregulation of piRNAs, rRNAs, and tsRNAs. Superinfection exclusion was robust throughout larval development and adulthood, providing protection against various OrV isolates. However, the protective effect of the initial infection diminished over time, suggesting different mechanisms of action and ultimately favoring the primary infecting virus. This phenomenon was dependent on a functional RNA interference pathway.

## INTRODUCTION

Superinfection exclusion (SIE) is a phenomenon in which a preexisting viral infection prevents a secondary one with the same or closely related viruses. It has been described for a range of viruses including human, plant and bacterial ones (Adams and Brown, 1985; Folimonova, 2012; Hofer et al., 1995; Michel et al., 2005; Webster et al., 2013). SIE can be mediated by a wide range of mechanisms, including blocking viral entry (Michel et al., 2005), RNA replication (Cherkashchenko et al., 2022), RNA interference (Gusachenko et al., 2021; Ratcliff et al., 1999), or unbalanced intracellular levels of viral components (Geib et al., 2003). SIE can be mediated by the host cell or by the primary virus, and may serve as a beneficial strategy by either minimizing the impact of infection or relaxing competition for cellular resources, respectively (Gusachenko et al., 2021; Hunter and Fusco, 2022; Ratcliff et al., 1999; Reitmayer et al., 2023).

The discovery of Orsay virus (OrV), the first natural virus of the nematode *Caenorhabditis elegans*, has allowed for the discovery of novel antiviral mechanisms, such as viral uridylation (Le Pen et al., 2018). OrV has a positive-sense, bi-segmented RNA genome of ∼6.3 kb which encodes four proteins: an RNA-dependent RNA polymerase (RdRP), a viral capsid protein (CP), the δ protein involved in nonlytic viral release, and a fusion of CP-δ that is incorporated into the virions as pentameric head fibers (Félix et al., 2011; Guo et al., 2020; Jiang et al., 2014; Yuan et al., 2018). OrV replicates in the intestinal epithelia of *C. elegans* (Félix et al., 2011; Franz et al., 2014), which consists of 20 single-layer cells surrounding the central lumen. Infection of OrV is restricted to the anterior intestinal cells and after cell egression transmits orofecally (Félix et al., 2011). The infection results in enlarged intestinal lumens and subcellular structural changes. However, the infection minimally affects animal life span, suggesting that the innate immune system is able to effectively restrain virus replication (Ashe et al., 2013). Amongst several innate immune mechanisms (*e.g*., physical barriers, antimicrobial compounds and induced responses), *C. elegans* uses RNA interference as the main defense mechanism against OrV (Castiglioni et al., 2024b; González and Félix, 2024; Zhou et al., 2024).

The infection of wild-type *C. elegans* by OrV is highly dynamic. We have previously reported four distinct infection phases throughout the larval development of wild-type animals using a highly concentrated viral stock and synchronous animal populations: (*i*) a pre-replication phase in which the viral load is indistinguishable from background levels, (*ii*) an exponential viral replication leading to a viral load peak, (*iii*) a phase of moderate viral load levels, and (*iv*) a phase of persistent residual infection, or resolution of the infection, in which the viral load came close to the initial background levels and OrV was only seen with a punctuated pattern in very few animals (Castiglioni et al., 2024b). These results suggested the possibility that OrV infection may be chronic and raised the possibility that the chronic infection would block secondary infections.

To tackle these possibilities, we have followed here the course of an infection throughout the lifespan of a synchronous wild-type animal population and described several successive peaks in viral load, suggesting that the infection is chronic. Moreover, we showed that the immune response mounted by the chronic infection is able to efficiently restrict viral replication of a secondary infection, indicative of superinfection exclusion. Analysis of the transcriptomic landscape at the time of re-infection showed a high number of differentially expressed genes, although the response had a lower magnitude than at the beginning of the infection. The small RNA (sRNA) landscape varied widely throughout the infection, with piwi-interacting RNAs (piRNAs), rRNAs, and tRNA-derived small RNAs (tsRNAs) being differentially expressed upon superinfection. Finally, we showed that the ability to control successive viral replications, but not the ability to ward off against different viral strains, decreases over time and is dependent on a functional RNA interference pathway.

## RESULTS

### Infection by OrV is chronic and peaks multiple times during adulthood

We have previously followed OrV infection during host larval development in synchronous populations and shown that at the end of larval development infection is either resolved or in a state of persistent residual infection, suggesting the potential for chronic infection. To address this possibility, we have followed the progression of the viral load in a synchronized population from hatching to senescence (Fig. 1A). We used the temperature-sensitive mutant *par-2(or640)*, which results in embryonic lethal defects at 25 °C, to study a single generation. This approach was chosen over other methods, such as the addition of Fluorodeoxyuridine (FUdR) to NGM plates to induce sterility, as FUdR has been shown to increase vertical vesicular stomatitis virus transmission in *C. elegans* (Gammon et al., 2017). We inoculated animals at the time of hatching, switched the temperature from 15 to 25 °C, and took samples at increasing hours post-inoculation (hpi) for quantitative RT-PCR (RT-qPCR). Viral load was determined by RT-qPCRs targeting RNA2 using the standard curve method for quantification. During larval development we observed a single peak (Fig. 1B), resembling what we reported previously at 20 °C (Castiglioni et al., 2024b) and a subsequent reduction of the viral load which occurred more drastically than at 20 °C. Wild-type animals were included as controls until 48 hours of post-hatching (hph), before the onset of egg-hatching, and showed similar viral load dynamics (Fig. 1B; Wilcoxon signed-rank test, *P* = 0.477). During adulthood we observed three subsequent peaks. The first peak was at 60 hpi, during the time of active egg laying, and had a higher magnitude than the peak at 12 hpi; the second peak was at 132 hpi, after the end of sexual reproduction, and had a comparable magnitude to the peak at 12 hpi; and the third was at 156 hpi and had the smallest magnitude. These successive viral load peaks indicate that the virus is able to replicate successfully hours after the host has managed to reduce its levels and that the infection at the end of larval development was a persistent residual infection.

**Figure 1.**
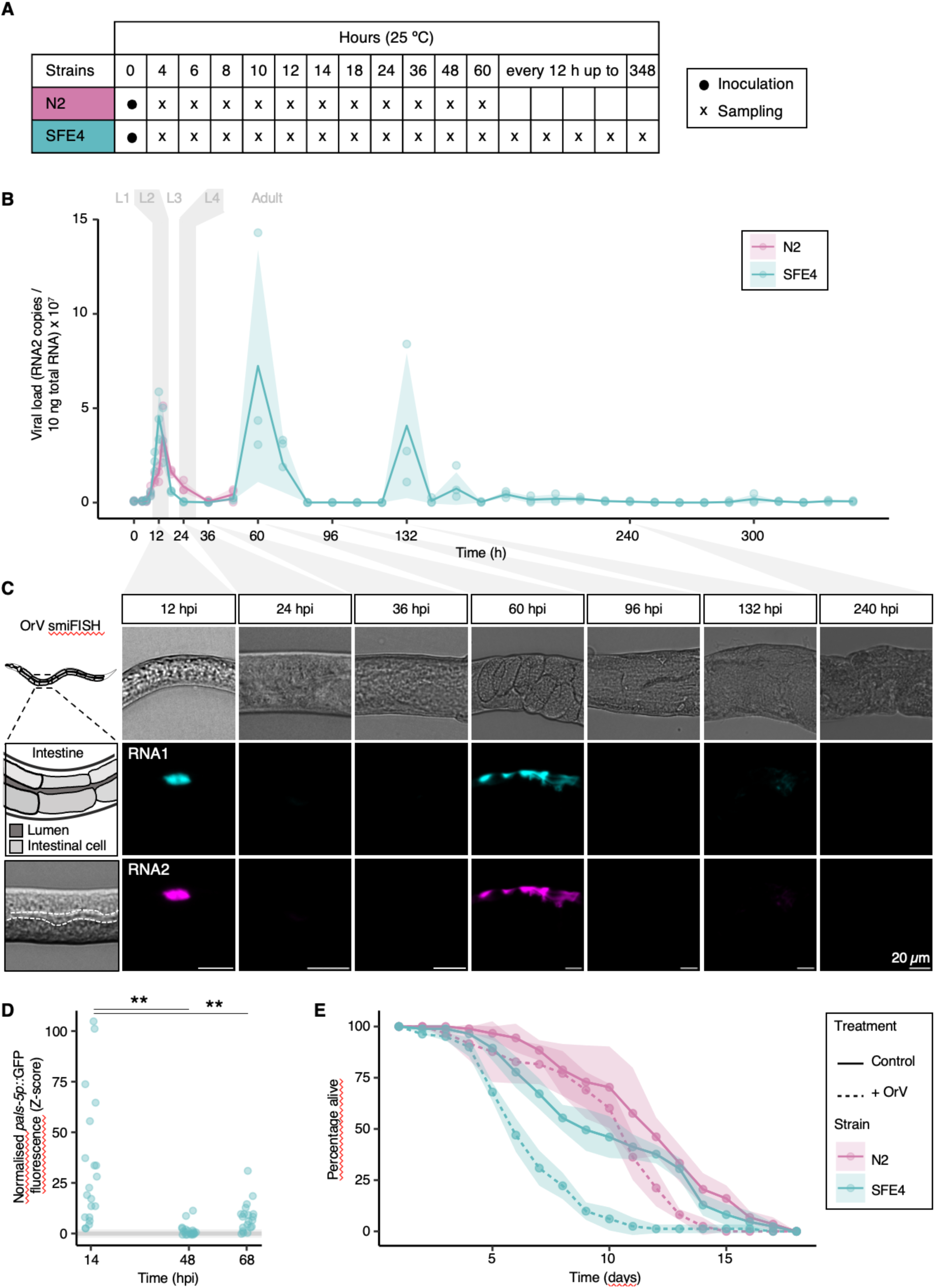
Infection by OrV is chronic and peaks multiple times during adulthood. (A) Experimental design. Synchronized recently hatched populations were inoculated with 2.8×10^9^ copies of OrV and grown at 25 °C. Samples were analyzed at the indicated timepoints by RT-qPCR or smiFISH. (B) Viral load determined over time by RT-qPCR. Viral load is defined as copies of RNA2/10 ng of total RNA. N2 in magenta, SFE4 in cyan. Solid lines and shading represent the mean ± 1 SD. *n* = 3 replicates per time point, each replicate consisting of a pool of 500 animals. The different developmental stages of the animal are represented in grey shading. (C) Representative smiFISH images of inoculated animals at the indicated timepoints. Top: DIC; middle: RNA1 in cyan; bottom: RNA2 in magenta. (D) *pals-5p::GFP* fluorescence intensity (*z*-score) of SFE12 animals at 14, 48 and 68 hpi normalized to the intensity of control animals. Horizontal grey lines indicate the range values of controls. *n* = 20 animals per time point. (E) Survival over time of N2 (magenta) and SFE4 (cyan) animals in control (solid line) and infected (dashed line) conditions. *n* = 3 replicates per condition, each replicate consisting of a pool of 30 animals.

To further characterize the chronic infection and confirm that the virus was amplifying again at late time-points, we inoculated animals at the time of hatching and took sequential samples for smiFISH analysis. RNA1 and RNA2 were labelled independently with 22 and 18 probes, respectively. We observed high smiFISH signal at 12 and 60 hpi, low signal at 132 hpi (Fig. 1C), and no signal at other time points, matching what we observed by RT-qPCR (Fig. 1B). These results were also consistent with the percentage of animals expressing high levels of the infection reporter *pals-5p::GFP* (Bakowski et al., 2014), which was 100% and 80% at 14 and 68 hpi, respectively, but only 30% at 48 hpi (Fig. 1D; Kruskal-Wallis test, χ^2^ = 43.799, 2 d.f., *P* < 0.001). The number of infected cells over time could not be assessed as sexual reproduction causes the germline to occupy most of the internal cavity of the animals, reducing the cellular resolution. We next investigated whether the survival of temperature-sensitive mutant *par-2(or640)* would be affected by OrV infection in a manner similar to that of wild-type animals. We followed the survival of infected and control wild-type and *par-2(or640)* animals. Firstly, we observed that *par-2(or640)* animals survived 14.38% less than wild-type animals (Fig. 1E; Kaplan-Meier survival regression: χ^2^ = 4.164, 1 d.f., *P* = 0.041). Secondly, more relevant, we found that infection resulted in significant reductions in the lifespan for both strains, with the detrimental effect being ∼2.6 times greater in *par-2(or640)* animals (Fig. 1E; *par-2(or640)*: 33.7% reduction, χ^2^ = 40.207, 1 d.f., *P* < 0.001; wild-type, 13.2% reduction χ^2^ = 15.666, 1 d.f., *P* < 0.001).

Altogether, our data shows that OrV infection becomes a chronic infection in late stages of life and that the virus is able to re-enter replicative cycles at late stages.

### *C. elegans* immunity, acquired during early larval stages, efficiently controls viral replication during early adulthood

We next investigated whether the decreases of viral load were due to an activation of the immune response which would suppress viral replication. If this was the case, animals exposed to a primary infection would efficiently contain a secondary infection whilst this response remained active. To test this, we inoculated wild-type animals that had been inoculated with OrV upon hatching, and would thus have an activated immune response, again at 38 hph, when the viral load from the primary infection was close to initial background levels (superinfected from hereon, SI.50h, Fig. 2A, B). We also inoculated animals at the time of hatching (early infection, EI.50h) and at 38 hph (late infection, LI.50h). We analyzed the viral load at 12, 38 and 50 hph by RT-qPCR (Fig. 2B) and RNA-seq (Fig. 2C). Consistent with our previous results (Castiglioni et al., 2024b), we observed a 96.5% reduction in the animals inoculated upon hatching than to those inoculated at 38 hph (Fig. 2B, EI.50h *vs* LI.50h; Bonferroni *post hoc* test, *P* < 0.001). The difference is likely due to the fact that animals inoculated upon hatching had time to mount a strong immune response against OrV whereas animals that had a single inoculation at 38 hph did not. We also observed that superinfected animals had very low viral loads, very similar to those of animals that had only been inoculated upon hatching or those that at 38 hpi had time to mount an efficient response and cleared off the virus (Fig. 2B, EI.50h *vs* SI.50h and EI.38h *vs* SI.50h; Bonferroni *post hoc* test, *P* = 1.000). The viral load in superinfected animals was 98.6% lower than in animals that had received the primary infection at 38 hph (Fig. 2B, SI.50h *vs* LI.50h; Bonferroni *post hoc* test, *P* < 0.001), suggesting that the exposure to a primary infection can limit viral replication on a secondary infection, indicative of SIE. The viral load at 12 hpi was not statistically significant, regardless of the time of infection (Fig. 2B, EI.12h *vs* LI.50h; Bonferroni *post hoc* test, *P* = 1.000).

**Figure 2.**
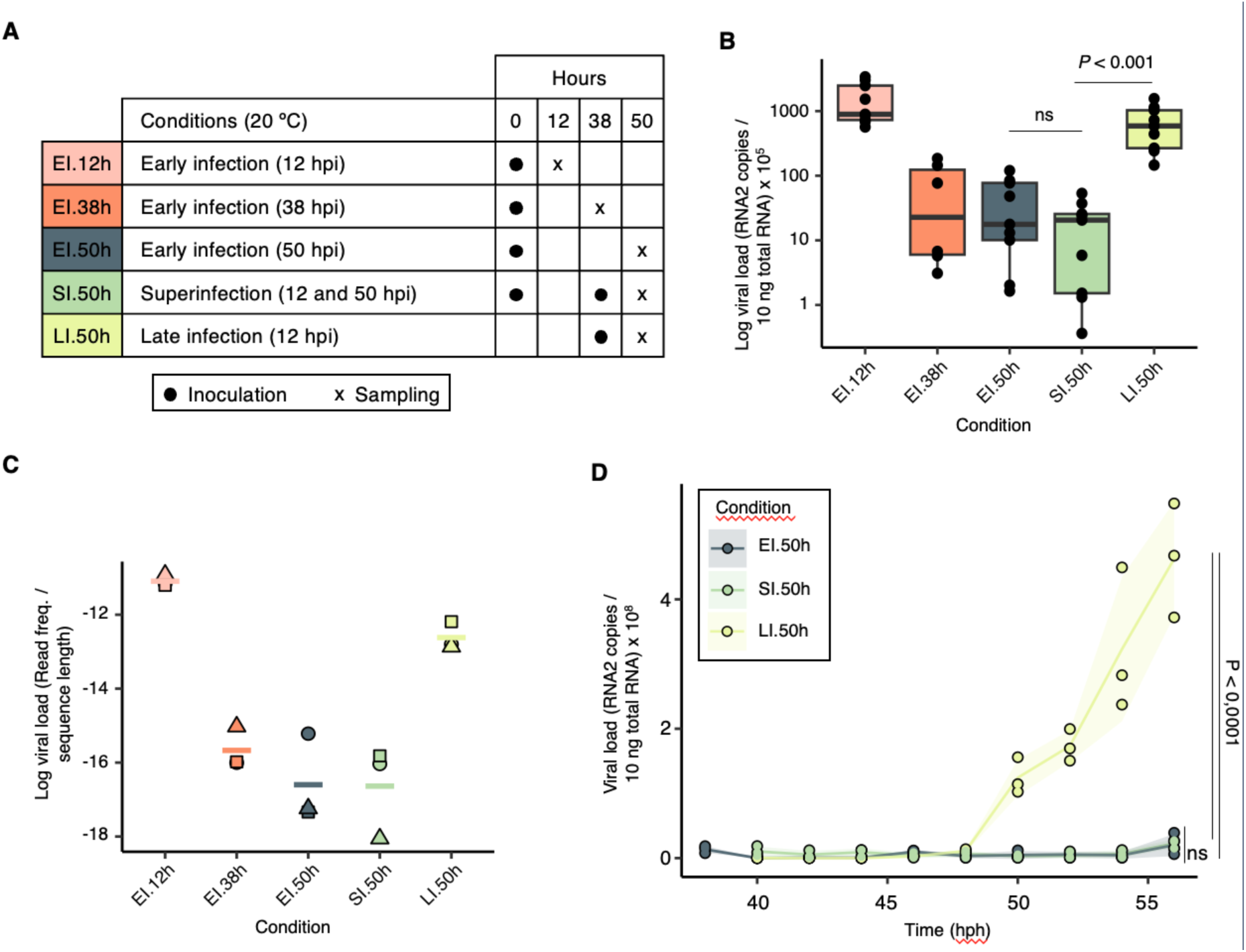
Exposure to a primary infection limits viral replication during a secondary infection. (A) Experimental design. Synchronized populations were inoculated with 2.8×10^9^ copies of OrV at the time of hatching or at 38 hph, grown at 20 °C and sampled at 12, 38 or 50 hph. Samples were analyzed by RT-qPCR. (B) Viral load (log RNA2 copies/10 ng total RNA) determined by RT-qPCR at 12 (light pink), 38 (orange) or 50 (dark green) hpi in populations infected upon hatching, upon superinfection (light green) or late infection (lime). *n* = 9 replicates per condition except for EI.38h, where *n* = 6, each replicate consisting of a pool of 500 animals. (C) Viral load (log read frequency/sequence length) determined by RNA-seq at 12 (light pink), 38 (orange) or 50 (dark green) hpi in populations infected upon hatching, upon superinfection (light green) or late infection (lime). *n* = 3 replicates (different symbols) per condition., each replicate consisting of a pool of 1200 animals. (D) Viral load (RNA2 copies/10 ng total RNA) over time by RT-qPCR upon 50 hpi (dark green), superinfection (light green) or late infection (lime). *n* = 3 replicates per condition, each replicate consisting of a pool of 500 animals.

As shown above and in our previous work (Castiglioni et al., 2024b), the viral load during OrV infection can be very dynamic. To confirm that the low viral load in superinfected animals was not due to a biased sampling, we followed the progression of the viral load from the second inoculation time-point (38 hph) to just before egg-hatching (56 hph) in the three conditions explained above in wild-type animals. We observed that the viral load remained at consistently low levels in the superinfection and in those that were inoculated upon hatching (Fig. 2D, EI.50h *vs* SI.50h; Bonferroni *post hoc* test *P* = 0.522), whilst it increased in the animals that were inoculated at 38 hph (Fig. 2C, SI.50h *vs* LI.50h; Bonferroni *post hoc* test, *P* < 0.001). This confirms that the immune response of animals that have been previously exposed to the virus efficiently contains subsequent viral replication events.

Altogether, these results suggest that superinfection exclusion controls OrV secondary infections.

### Superinfection results in a more specific transcriptomic defense response

To better understand the ability to control viral replication in superinfected animals, we next sought to evaluate the extent in which the transcriptomic response to superinfection is different from a primary infection. For this purpose, we inoculated animals at the time of hatching and took samples at 12 (EI.12h), 38 (EI.38h) and 50 hpi (EI.50h), at the time of hatching and at 38 hpi to analyze at 50 hph (SI.50h), and at 38 hph to analyze at 50 hph (LI.50h). Non-inoculated controls were taken at 12, 38 and 50 hph (Fig. 3A). Differential expression analysis (DEA) by time point showed that the highest number of differentially expressed genes (DEGs) was at 38 hpi, with a predominance of down-regulated genes (Fig. 3B, EI.38h), but most of these genes had a small magnitude of change (|log_2_*FC*| < 2, Fig. 3C). In turn, the magnitude of the transcriptomic response was largest during early infection (EI.12h and LI.50h), with most DEGs having log_2_*FC* > 2 and a predominance of up-regulated genes. The transcriptomic response was diminished both in magnitude and quantity of DEGs at 50 hpi. Importantly, superinfection resulted in a smaller transcriptomic response, both in magnitude and quantity of DEGs, and most similar to EI.50h (Fig. 3C), suggesting that a primary viral infection diminishes the activation of a transcriptomic response upon the subsequent inoculation.

**Figure 3.**
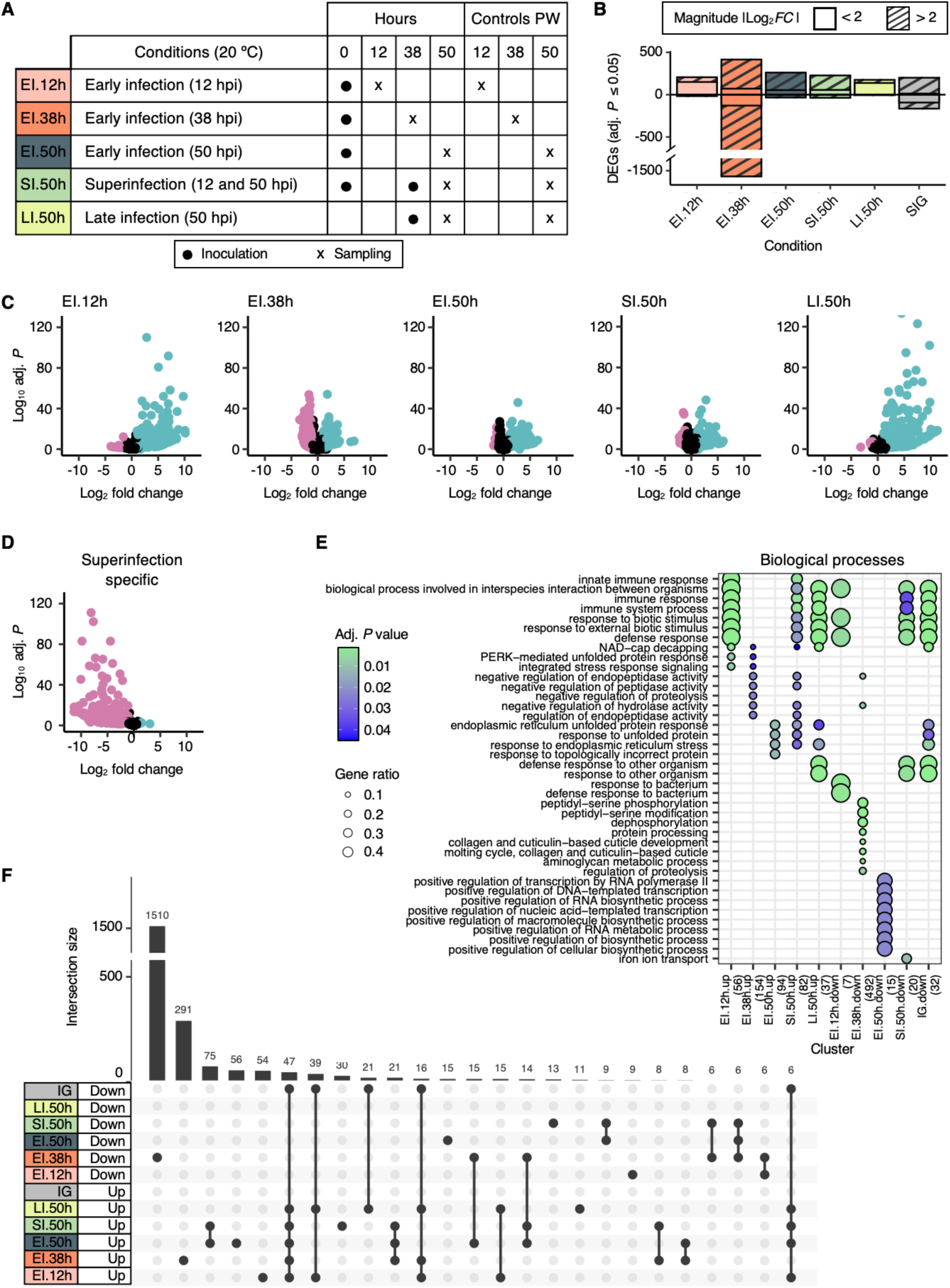
The transcriptomic response to superinfection is small, primed and specific. (A) Experimental design. Synchronized populations were inoculated with 2.8×10^9^ copies of OrV at the time of hatching or at 38 hph, grown at 20 °C and sampled at 12, 38 or 50 hph. Synchronized control populations, used for the pairwise comparison, were grown at 20 °C and sampled at 12, 38 or 50 hph. Samples were analyzed by RNA-seq. (B) Number of DEGs with an adjusted *P* ≤ 0.05 and a |log_2_*FC*| ≥ 1 after the pairwise comparison of the conditions introduced in (A) and superinfection immunity genes (SIG). Solid pattern indicates 2 > |log_2_*FC*| ≥ 1 and striped pattern indicates |log_2_*FC*| ≥ 2. (C) Differentially expressed genes at each condition. Cyan and magenta represent up-regulated and down-regulated genes (adjusted *P* < 0.05 and |log_2_*FC*| ≥ 1), respectively. (D) Differentially expressed superinfection specific genes. Cyan and magenta represent up-regulated and down-regulated genes (adjusted *P* < 0.05 and |log_2_*FC*| ≥ 1), respectively. (E) Functional enrichment of biological processes of the conditions introduced in (A) and superinfection immunity genes (SIG). Only significant terms are shown (adjusted *P* ≤ 0.05). (F) Upset plot showing the intersections between the DEGs of each condition. Only intersections with at least six genes are shown. The points indicate the sets that are counted in the vertical bars.

When looking at the GO-terms associated with each response, we observed the strongest enrichment in immune related processes within the DEGs from early infection (EI.12h and LI.50h, Fig. 3E). At 38 and 50 hpi we observed a predominance in metabolism and stress related GO-terms. Even though the size and magnitude of the transcriptomic response upon superinfection were small, many of the GO-terms associated with superinfection were related to the immune response, indicating that the primary infection does not prevent reactivation of transcriptomic defense responses. Moreover, many of the GO-terms associated with superinfection were observed at 38 hpi but not at 12 or 50 hpi, suggesting that the primary infection primes some genes to react earlier. The molecular function GO-terms associated with superinfection were most similar to a late response (Suppl. Fig. S1A, EI.50h), whilst the cellular component GO-terms associated with superinfection fell within the same categories as with an early infection (Suppl. Fig. S1B, SI.50h (up), and EI.12h), or a late response (SI.50h (down) and EI.50h). The enrichment in KEGG pathways upon superinfection was very similar to either primary early infection (Suppl. Fig. S1C, EI.12h or LI.50h).

We then concentrated on the superinfection-specific transcriptomic response, *i.e*., responses that were observed upon superinfection but not upon a primary infection (either early or late in the animal’s development) (Fig. 3B, D). Within the transcriptomic responses, we observed that superinfection resulted in specific upregulation of only seven genes: *F52F10.2*, *fbxa-84*, *hrg-1*, *oac-6*, *T06C12.14*, *ugt-44*, and *Y75B8A.28*. Whilst *fbxa-84*, *ugt-44* and *Y75B8A.28* have been previously shown to be differentially expressed by OrV at some point of the larval development (Castiglioni et al., 2024b), the role of these seven genes in OrV infection is unknown. A much greater number of genes (158) was found to be downregulated upon superinfection *vs* a primary infection. The downregulated genes were related to defense, metabolism and developmental processes (Fig. 3F), and further suggests that a primary infection can trigger immune responses that control a secondary infection in part due to a more efficient transcriptomic response.

### Superinfection alters the sRNA landscape

We next sought to describe the sRNA differentially responding to superinfection. In contrast to the transcriptomic response, the endogenous sRNA response had a higher magnitude and number of differentially expressed sequences (DES) at 38 hpi (Fig. 4A, B). Most of the DES at 38 hpi were down-regulated 26G RNAs, primary endogenous sRNAs of the RNAi pathway produced by the enhanced RNAi (ERI) complex (Fig. 4C, D), suggesting that the endogenous RNAi pathway is being repressed at 38 hpi. The highest number of upregulated miRNAs, involved in RNA silencing and post-transcriptional gene silencing, was also observed at 38 hpi (Fig. 4E, F). Interestingly, some of the DEGs at this time have been reported to be targets of up to five differentially expressed miRNAs at this timepoint, suggesting redundancy and robustness in the regulatory network (Suppl. Fig. S2). Early infection of young animals resulted in less upregulated miRNAs, but a bigger magnitude of change), suggesting a role for miRNAs during early infection of young animals. Interestingly, a downregulation of miRNAs was observed at late infection times and superinfection, and a very small upregulation upon early infection of older animals (LI.50h), hinting at developmental-dependent differences in the miRNA responses.

**Figure 4.**
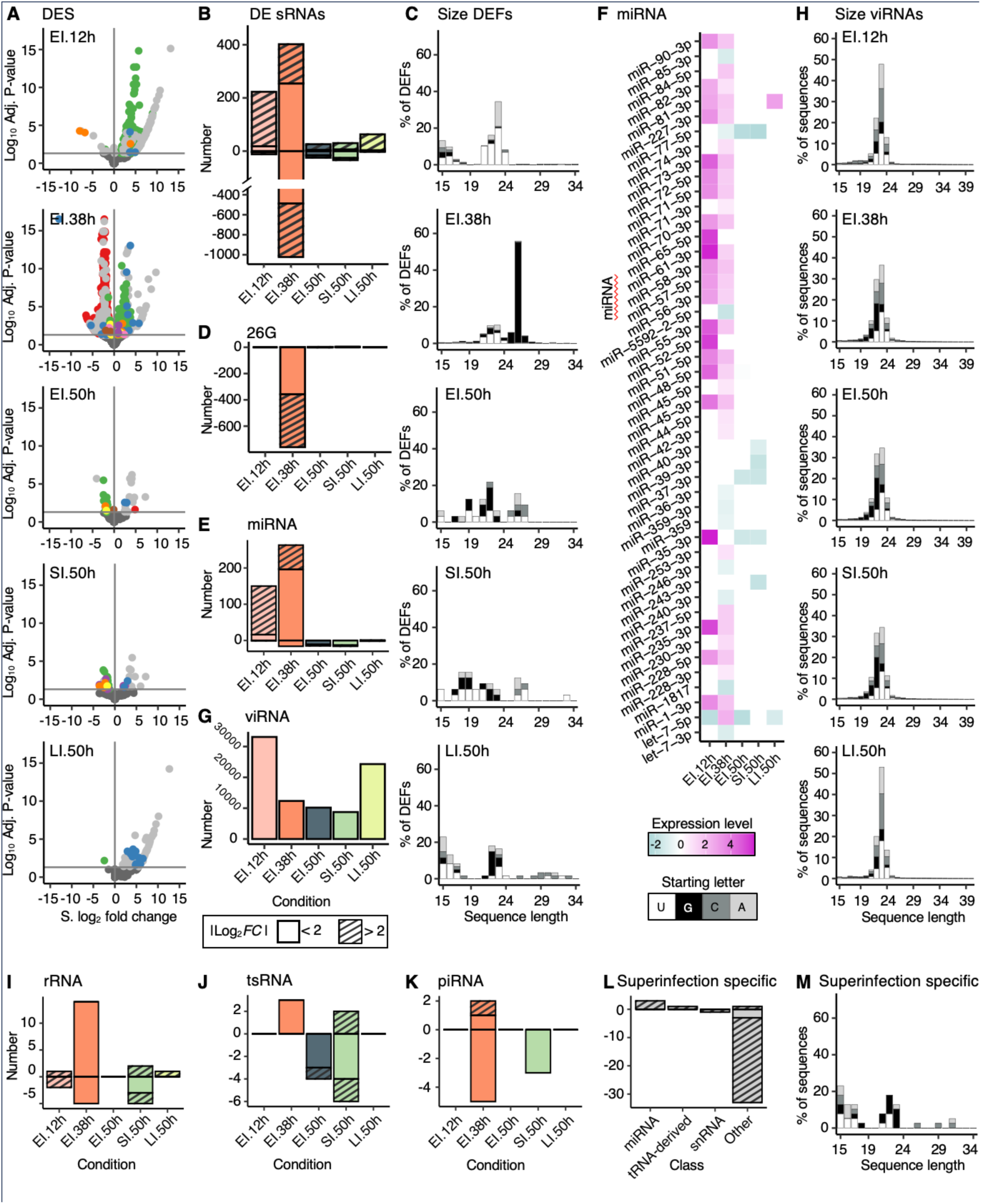
Superinfection alters the sRNA landscape. (A) Differentially expressed fragments (adjusted *P* < 0.05) in each condition. RNA class are colored as follows: 26G-RNAs (red), miRNAs (green), tsRNAs (purple), small rRNAs (orange), snRNAs (yellow), snoRNAs (brown), piRNAs (pink), other 23 nt RNAs (blue), other unclassified small RNAs (light grey) and non-significant sRNAs (dark grey). (B) Number of upregulated and downregulated fragments (adjusted *P* < 0.05) in each condition. Solid pattern indicates 2 > |log_2_*FC*| ≥ 1 and striped pattern indicates |log_2_*FC*| ≥ 2. (C) Percentage of differentially expressed fragments of each sequence length aligning to *C. elegans* genome in each condition. (D) 26G-RNAs in each condition. (E) Number of upregulated and downregulated miRNAs in each condition. (F) Differential expression profile of the miRNAs with adjusted *P* < 0.05. (G) Number of upregulated and downregulated viRNAs in each condition. (H) Percentage of fragments of each sequence length aligning to OrV genome in each condition. (I) Number of upregulated and downregulated small rRNAs in each condition. (J) Number of upregulated and downregulated tsRNAs in each condition. (K) Number of upregulated and downregulated piRNAs in each condition. (L) Number of superinfection specific sRNAs resulting from comparing superinfection with early and late primary infections. (M) Percentage of differentially expressed fragments of each sequence length aligning to *C. elegans* genome resulting from comparing superinfection with early and late primary infections (superinfection specific).

Twenty-three nt sRNAs, primary short interfering RNAs (siRNAs) produced by DCR-1, were the most abundant sRNAs across all timepoints in control and infected animals (Suppl. Fig. S3A). Twenty-three nt sRNAs mapping against the virus genome were predominantly observed during early infection (EI.12h and LI.50h, Fig. 4G, H), correlating with the observed viral load at these time points (Fig. 2B). The lowest small viral RNA (viRNA) counts were observed upon superinfection (Fig. 4G), suggesting that a primary viral infection also prevents the production of viral siRNAs upon superinfection.

Apart from classification of the DES into 26G sRNAs, miRNAs and viRNAs, classification of the DES into further classes showed distinct enrichments in the different conditions. Early infection later in life and early infection at 50 hpi (LI.50h and EI.50h) showed no predominant up- or downregulation and no enrichment of any particular sRNA length (Fig. 4B, C). Of note, superinfection resulted in the misregulation of rRNA, tsRNAs, and downregulation of piRNAs (Fig. 4I, J, K respectively). These three sRNA classes had more DES upon superinfection than upon any primary infection (EI.50h or LI.50h), with tsRNAs showing the highest number of DES upon superinfection than in any other condition (Fig. 4J). Moreover, we observed downregulation of snoRNAs at 38 and 50 hpi, but not upon superinfection (Suppl. Fig. S3B) and downregulation of snRNA mostly at 38 hpi (Suppl. Fig. S3C).

When looking at the superinfection specific sRNA response, the response that was observed upon superinfection but not upon a primary early or late infection, we identified upregulation of miRNA *let-7*, tsRNA *5p-tRF*, snRNA *sls-1.1*. Additionally, we noted the downregulation of several other sRNAs, among which are *aex-5*, which regulates defecation (Mahoney et al., 2008), *unc-54*, which physically interacts with the δ protein of OrV (Fan, 2018), and *vet-2*, which interacts with autophagosome protein *lgg-1*, among others (Fig. 4L, M, Suppl. Table S3).

Altogether, these results indicate that the sRNA landscape is very dynamic. We observed a down-regulation of the endogenous RNAi pathway at 38 hpi, and no indications of differential expression at other time points. In turn, the antiviral RNAi response seems to correlate to the viral load. miRNAs are upregulated at early infection time points during early developmental stages, but not during later developmental stages. Finally, piRNA, rRNAs and tsRNA are misregulated upon superinfection.

### Functioning of SIE and the role of RNAi

We have shown above that the immune response mounted by a chronic infection is able to control viral replication during early adulthood. However, at the time of the secondary infection (38 hph) there was a big downregulation of transcripts and 26G RNAs (Fig. 3B, Fig. 4B). To determine whether these responses are necessary to contain viral replication, we delayed the time of secondary infection to 50 hph (SI.62h) and compared the viral load to that of animals inoculated at the time of hatching (EI.62h) and at 50 hph (LI.62h). In all cases we analyzed the viral load at 62 hph by RT-qPCR (Fig. 5A). We observed that SI.62h and EI.62h had comparable viral loads (Fig. 5B; Bonferroni *post hoc* test, *P* = 1.000), which were over three orders of magnitude lower than the viral load of LI.62h (Fig. 5B; Bonferroni *post hoc* test, *P* < 0.001), indicating that the primary infection at the time of hatching can contain viral replication upon a secondary infection during adulthood.

**Figure 5.**
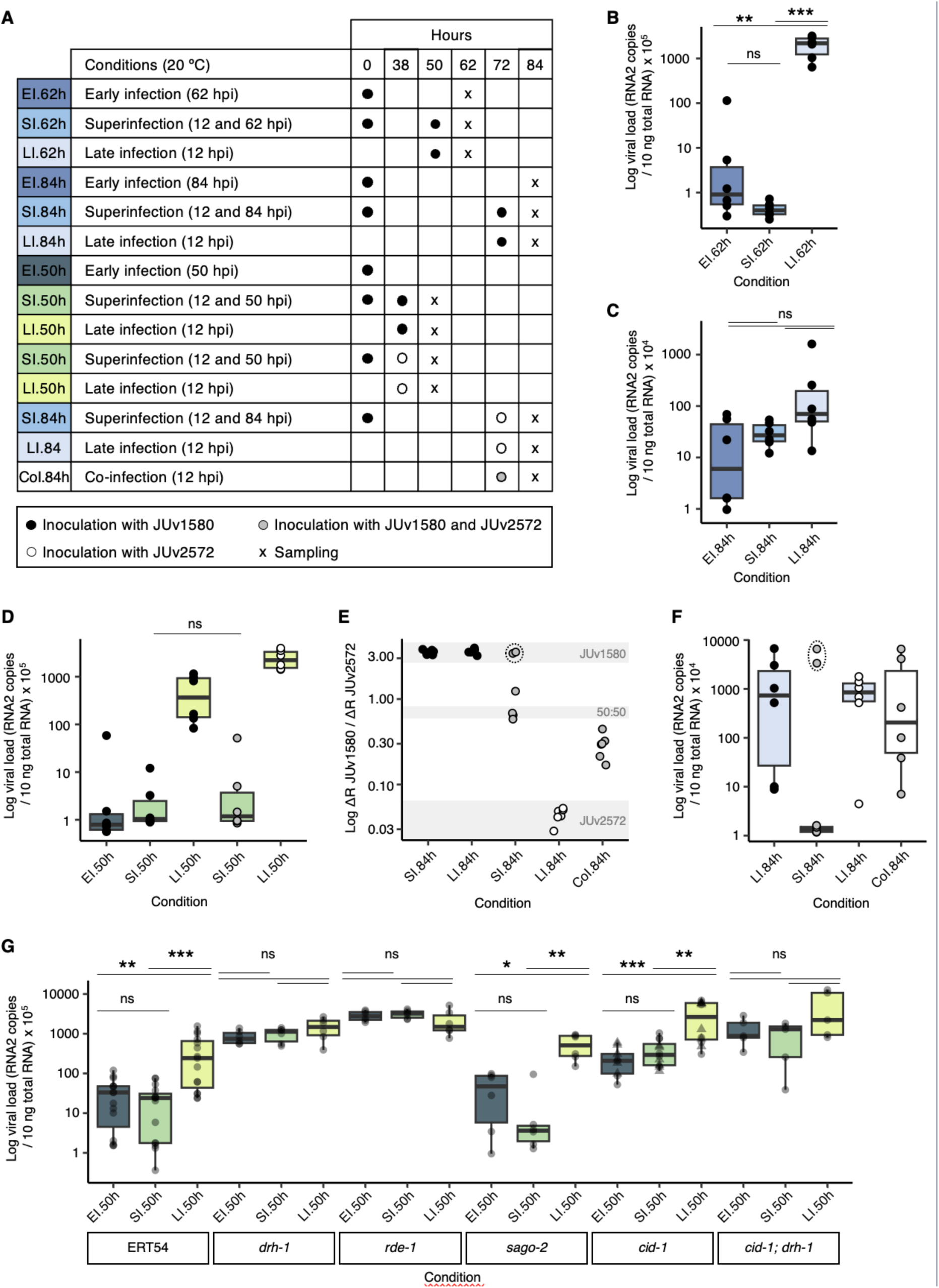
Characterization of the temporal decaying immunity and superinfection exclusion and contribution of RNAi to SIE. (A) Experimental design. Synchronized populations were inoculated with 2.8×10^9^ copies of OrV JUv1580 or JUv2572 at the time of hatching, 38, 50 or 72 hph, grown at 20 °C and sampled at 50, 62 or 84 hph. Samples were analyzed by RT-qPCR or TaqMan qPCR. (B) Log-viral load (RNA2 copies/10 ng total RNA) determined by RT-qPCR at 62 hpi in populations inoculated upon hatching (dark blue), superinfection (blue) or late infection (light blue). *n* =6 replicates per condition. (C) Log-viral load (RNA2 copies/10 ng total RNA) determined by RT-qPCR at 84 hpi in populations inoculated upon hatching (dark blue), superinfection (blue) or late infection (light blue). *n* = 6 replicates per condition. (D) Log-viral load (RNA2 copies/10 ng total RNA) determined by RT-qPCR at 50 hpi in populations inoculated upon hatching (dark green), superinfection (green) or late infection (lime) with JUv1580 or JUv2572. Black data points indicate samples were inoculated with JUv1580, grey indicates samples were inoculated with both JUv1580 and JUv2572 and white indicates they were inoculated with JUv2572. *n* = 6 replicates per condition. (E) ΔR JUv1580/ΔR JUv2572 determined by TaqMan qPCR of RNA1 at 84 hpi in superinfected or late infected populations. Black data points indicate samples were inoculated with JUv1580, grey indicates samples were inoculated with both JUv1580 and JUv2572 and white indicates they were inoculated with JUv2572. Horizontal grey shading indicates values obtained for populations inoculated with JUv1580 (top), JUv2572 (bottom) or a 50:50 mixture of both (middle). *n* = 6 replicates per condition. (F) Log-viral load (RNA2 copies/10 ng total RNA) determined by RT-qPCR at 84 hpi in superinfected populations (blue), late infected (light blue) or coinfected populations (white). Black data points indicate samples were inoculated with JUv1580, grey indicates samples were inoculated with both JUv1580 and JUv2572 and white indicates they were inoculated with JUv2572. *n* = 6 replicates per condition. (E) Log-viral load (RNA2 copies/10 ng total RNA) determined by RT-qPCR at 50 hpi in populations of N2, *drh-1(ok3495)*, *rde-1(ne219)*, *sago-2(tm894)*, *cid-1(tm936)* and *cid-1(tm1021),* represented together with circles and triangles, respectively, and double mutant *cid-1(tm936);drh-1(ok3495)* inoculated upon hatching (dark green), superinfection (green) or late infection (lime). *n* = 6 replicates per condition except for N2, where *n* = 15. Bonferroni’s *post hoc* test was used for all pairwise comparisons.

Given that viral replication can be re-activated in adult worms around the time of maximum egg laying (∼60 hph at 25 °C, ∼84 hph at 20 °C), we tested whether the immune response of older animals would also restrict viral replication when superinfected. For this purpose, we inoculated animals at the time of hatching (EI.84h), at 72 hph (LI.84h) and both at the time of hatching and at 72 hph (SI.84h), and measured the viral load at 84 hph, around the time of maximum egg laying (Fig. 5A). In qualitative agreement with the results described in the previous paragraph, the viral load of SI.84h and EI.84h animals was undistinguishable (Fig. 5C; Bonferroni *post hoc* test, *P* = 0.493). However, the viral load of LI.84h animals was significantly larger than observed in EI.84h animals (Fig. 5C; Bonferroni *post hoc* test, *P* = 0.027) but not from SI.84h animals (Fig. 5C; Bonferroni *post hoc* test, *P* = 0.439). In this cellular context, superinfection resulted in an intermediate phenotype, in which the viral load was bigger than upon a late infection but smaller than a primary infection, suggesting that the protective effect, whilst still there, is diminished with age.

We next sought to test whether the SIE would also prevent a secondary infection with a different OrV isolate, JUv2572 (Castiglioni et al., 2024a; Frézal et al., 2019). We inoculated wild-type animals that had been primarily exposed to isolate JUv1580 at 38 hph with either JUv1580 or JUv2572 (SI.50h). We also inoculated animals at the time of hatching with JUv1580 (EI.50h) and at 38 hph with either JUv1580 or JUv2572 (LI.50h). In all cases, we analyzed the viral load at 50 hph by RT-qPCR (Fig. 5A). A primary infection with JUv1580 limited viral replication upon superinfection with JUv1580 and JUv2572 to a comparable extent (Fig. 5D. JUv1580 SI.50h *vs* JUv2572 SI.50h; Bonferroni *post hoc* test, *P* = 1.000) indicating that SIE can prevent secondary infections with different isolates.

Given that the protective effect of a primary infection was weaker at later timepoints, we subsequently asked whether the diminished effect was due to loss of SIE capacity or due to the virus overcoming the intracellular defenses in older animals. We inoculated wild-type animals that had been primarily exposed to isolate JUv1580 (SI.84h), as well as *naïve* animals (LI.84h), at 72 hph with either JUv1580 or JUv2572. For comparison, we also co-inoculated *naïve* animals at 72 hph with equimolar concentration of JUv1580 and JUv2572. We analyzed the proportion of JUv1580 and JUv2572 in these samples, as well as the viral load by RT-qPCR, at 84 hph (Fig. 5A). Samples inoculated with JUv1580 or JUv2572 had high or low accumulation values, respectively, whilst the equimolar mixture of JUv1580 and JUv2572 lay in the middle as expected (Fig. 5E). Superinfection with JUv2572 showed two distinct populations, with four out of six samples close to the equimolar mixture and the other two being consistent with the only presence of JUv1580. These values allow us to compute the fitness of JUv1580 relative to JUv2572 as in Carrasco et al. (2007), rendering a value of 1.076 ±0.075. Interestingly, no sample showed more JUv2572 than JUv1580, suggesting that SIE is still active in older animals. By contrast, when both viruses were coinfected (CoI.84h), the outcome was radically different, resulting in significantly more JUv2572 accumulation (JUv1580 fitness relative to JUv2572 of 0.930 ±0.025; *z*-test *P* < 0.001), indicating that JUv2572 has a higher replicative capacity than JUv1580 in a period of 12 hpi in absence of SIE induced by JUv1580. When analyzing viral loads, we observed similar values in animals that had received a single inoculation with either JUv1580 or JUv2572 or had been coinfected with both viruses (Fig. 5F; Bonferroni *post hoc* test, *P* ≥ 0.919). Superinfection with JUv2572 showed the same two distinct populations, with the four samples that had close to equimolar mixture showing very low viral loads, and the two samples with JUv1580 amplification (encircled with dotted line) showing high viral loads. These results suggest that SIE is still active in older animals and that increased viral loads at later timepoints are either due to a weakened immune system or viral dynamics, but not due to new viral uptake.

Finally, we sought to assess the contribution of two antiviral pathways, the RNA interference (RNAi) and viral uridylation pathways, to the acquired immunity and its ability to limit viral replication during early adulthood. For this purpose, we compared the response of superinfected animals to that of animals which had been inoculated at 38 hph and at the time of hatching (Fig. 5A). Mutants of *drh-1*, which is involved in viral recognition during the RNAi pathway; *rde-1*, which is a primary Argonaute protein (Ago) downstream of *drh-1*; *sago-2*, which is a secondary Ago downstream of *rde-1*; and *cid-1*, the viral uridyltransferase, were analyzed. Viral loads were analyzed at 50 hph by RT-qPCR. We observed that loss of *sago-2* or *cid-1* did not affect the capability of mounting an immune response which would contain viral replication upon superinfection, similar to what we observed in wild-type animals (Fig. 5G; Bonferroni *post hoc* test, *P* ≤ 0.008 in both cases). In contrast, loss of RNAi components *drh-1* and *rde-1* failed to mount an immune response in superinfected animals which would contain viral replication (Fig. 5G; Bonferroni *post hoc* test, *P* ≥ 0.199 in both cases), suggesting that the RNAi response is necessary for SIE. Consistent with this, the viral load at 50 hpi upon loss of *drh-1* and *rde-1* was not distinguishable from the viral load at 12 hpi, suggesting that these genes are important for containing viral replication at late infection stages (Mann-Whitney *U* test, *P* ≥ 0.132 in both cases).

To further explore the interactions between the RNAi and viral uridylation pathways, we also assessed the effect of inactivation of both pathways simultaneously (double mutant *cid-1; drh-1*), and observed similar results as upon loss of RNAi components *drh-1* and *rde-1*: a failure to mount an immune response in superinfected animals which would restrict viral replication, resulting in undistinguishable viral loads at 12 and 50 hpi (Mann-Whitney *U* test, *P* = 0.786). The viral load at 50 hpi was significantly higher upon simultaneous loss of *cid-1* and *drh-1* than upon loss of *cid-1* (Fig. 5G; Mann-Whitney *U* test, *P =* 0.004), but not higher than upon loss of *drh-1* individually (Fig. 5G; Mann-Whitney *U* test, *P* = 0.537), suggesting that loss of RNAi is primarily responsible for the phenotype observed upon inactivation of both pathways.

Altogether, the results from this section indicate that the ability to repress viral replication upon superinfection persists until adulthood but wanes over time. Moreover, SIE can prevent a secondary infection with a different strain of OrV even in older animals, indicating that viral replication in older animals is due to reactivation of the primary infection. Finally, our results suggest that the RNAi response is essential for the superinfection response.

## DISCUSSION

Herein, we have followed the progression of a viral infection throughout the whole lifespan of *C. elegans* and characterized the superinfection exclusion (SIE) response of the animal to OrV. We observed that OrV infection becomes chronic in *C. elegans*, with several phases of viral replication followed by periods in which the virus remain at low background levels, reminiscent of other persistent infections, *e.g*., herpes simplex virus-2 infection in humans (Schiffer et al., 2018). Moreover, exposure to a primary infection limited viral replication during a secondary infection of the same or a closely related OrV strain, indicative of SIE. The protective effect granted by a primary infection diminished around the time that animals reached their peak reproductive stage, but SIE remained active, suggesting distinct protective mechanisms. Interestingly, we found that SIE was dependent on a functional RNAi pathway.

### Superinfection exclusion in *C. elegans*

It remains unclear though whether SIE is a virus-mediated mechanisms that provides a competitive advantage to the primary virus or a consequence of the animal’s immune system.

Mechanisms of SIE are diverse and have not been determined in all cases, but all mechanisms described so far depend on direct interaction of products of the primary infection with the superinfecting virus and can be distinguished between mechanisms that prevent viral entry or fusion and post-entry mechanisms (Cwick et al., 2022; DaPalma et al., 2010). Blocking cellular entry or early steps in the viral cycle, prior to the detection of the virus by the RNAi machinery, has been reported for some *Caenorhabditis briggsae* strains and *Nodaviridae* (Alkan et al., 2024). *C. elegans* could promote SIE by several mechanisms: (*i*) RNAi, wherein sequences homologous to the viral RNA will be present in the cell upon entry of the secondary infecting virus; (*ii*) expression of different antiviral genes during the primary infection which can confer protection to a secondary infection, such as collagens, which are upregulated upon infection and prevent OrV entry (Castiglioni et al., 2024b; Zhou et al., 2024), or (*iii*) through other unidentified mechanisms, such as by priming defense genes to respond more efficiently upon a secondary infection, as shown in plants (Corrêa et al., 2024).

On the one hand, several results suggest that SIE is based in part in a more efficient transcriptomic response: (*i*) superinfection resulted in a smaller transcriptional response regarding number of DEGs than a chronic infection and a weaker response than a late primary infection regarding the magnitude of change of the DEGs, (*ii*) a transcriptional downregulation of immune-related genes was observed when comparing a superinfection to both primary infections, but this did not allow for viral replication, and (*iii*) many of the GO-terms observed early upon superinfection were only observed at a late time point of a primary infection, suggesting that the primary infection primes some genes to react earlier. Superinfection also altered the sRNA landscape in comparison to either primary infection, with a misregulation of tsRNAs and small rRNAs and downregulation of piRNAs, all of which have various links to stress and viral infections (Cao et al., 2014; Jiang et al., 2024; Li et al., 2018; Zhou et al., 2017). The functional relevance of these alterations remains to be determined.

On the other hand, SIE seems to depend on a functional RNAi pathway. SIE has been previously proposed to be a consequence of cross-reactive RNAi, with viruses with lower genetic divergence showing more pronounced SIE (Gusachenko et al., 2021; Ratcliff et al., 1999). SIE of citrus tristeza virus occurs only between isolates of the same strain, usually showing less than 5% genomic divergence, but not between isolates of different strains (Folimonova et al., 2009). We have tested here SIE with the two only isolates of OrV identified so far, JUv1580 and JUv2572, which have a genomic divergence of 2.3% (Castiglioni et al., 2024a). Use of RNAi mutants indicated that the RNAi response is essential for SIE in *C. elegans*. Moreover, a decrease in the efficiency of RNAi over time could explain the subsequent peaks in viral load in aging animals, as the RNAi response in aging *C. elegans* has been shown to decay over time (Knudsen-Palmer et al., 2024). Whilst in our study a primary infection could limit viral replication during the first 2-3 dpi, with a weakened effect at 3-4 dpi, previous studies have observed that the RNAi response to an exogenous stimuli can last up to eight days (Knudsen-Palmer et al., 2024). The shortened duration of the response we observed could be due to the different tissues or targets assayed, as Knudsen-Palmer et al. (2024) tested the response against an endogenous gene in muscle cells and we are studying the response to an intestinal virus, which could have evolved some yet to be described mechanism to avoid RNAi. Differences in the mechanics of the RNAi response to endogenous targets and OrV are numerous, most strikingly is perhaps the absence of an inherited transgenerational response to OrV (Ashe et al., 2015), whilst RNAi against endogenous targets can last several generations (Vastenhouw et al., 2006). Discovery of more isolates of OrV, and particularly of isolates with a higher genomic divergence, will allow for a more comprehensive analysis of the role of RNAi in SIE. However, a loss of functionality in the RNAi pathway alone fails to explain why a different viral strain in older animals does not replicate, and opens the possibility that a combination of mechanisms is responsible for SIE. These results are in agreement to what was observed for some other *Nodavirus,* which have been reported to block cellular entry or early steps in the viral cycle prior to the detection of the virus by the RNAi machinery (Alkan et al., 2024).

While SIE was mainly beneficial for the primary infecting virus, even when the primary infecting virus had a lower replicative capacity than the secondary infecting virus. Given that some viruses capable of SIE, such bacteriophages, are capable of overtaking faster growing populations incapable of SIE even if this ability comes at a significant cost to their growth rate (Hunter and Fusco, 2022), it would be interesting to know whether other OrV strains with higher replicative capacity are capable of SIE and whether experimental evolution under conditions which favor SIE result in decreased growth rates. Indeed, genetic drift is minimized when superinfection occurs, facilitating the fixation of beneficial mutations and the removal of deleterious ones (Hunter and Fusco, 2022), resulting in populations which are less capable of adapting to changes in the environment. The analysis of the diversity within superinfected populations and within populations evolved under superinfection conditions, and their fitness after selection, will allow us to assess the impact of SIE in the long-term evolution of OrV and could help us understand whether this mechanism is mediated by the virus.

In conclusion, SIE of OrV in *C. elegans* seems to be mediated by a more efficient transcriptomic response in combination with a primed RNAi pathway. Even though the primary infecting virus is greatly benefited by SIE, it remains uncertain whether the virus is partly responsible for this mechanism. The effect of SIE on OrV populations evolvability also remains to be determined. Discovery of more isolates of OrV, and particularly of isolates with higher genomic divergence, will greatly aid in answering these open questions.

### Insights into OrV infection dynamics

Several insights derived from this study are worth mentioning. Firstly, in a previous study we described a viral peak followed by a viral load decrease, a phase of constant viral load and then a decrease to background levels, all within larval development at 20 °C (Castiglioni et al., 2024b). Here we reported one viral load peak followed by a sharp decrease to background levels within the larval development at 25 °C, with no intermediate phase of constant viral load. Development of *C. elegans* at 20 and 25 °C is widely different (Altun and Hall, 2009; Begasse et al., 2015), thus differences that arise because of temperature are more likely due to the host. However, the viral load peak was observed at 12 hpi at both temperatures, suggesting that the phase of exponential viral replication is determined by viral dynamics rather than the host. In turn, the following phases of reduction and maintenance of stable viral load levels that we observed at 20 °C do not take place at 25 °C, suggesting that host factors are more relevant at these latter phases. Whilst a mild temperature increase from 20 to 25 °C is not supposed to have heat-shock protective effects reported against OrV (Castiglioni and Elena, 2024; Huang et al., 2021), temperatures of 25 °C can have a significant impact in sRNA populations (Conine et al., 2010; Conine et al., 2013; Gu et al., 2009; Seroussi et al., 2023), which could in turn be impacting the defense response against OrV.

Secondly, while inoculation at time of hatching (early) and at the end of larval (late) development resulted in very similar transcriptomic responses, the time of inoculation had a big effect on the miRNAs that were differentially expressed upon infection: early inoculation resulted in a strong upregulation of multiple miRNAs, while upregulation at late larval development only affected two miRNAs. Superinfection, in turn, resulted in downregulation of several miRNAs. miRNAs modulate several cellular processes through post-transcriptional gene regulation and several RNA viruses have been reported to interact directly with cellular miRNAs and/or to use these miRNAs to augment their replication potential. miRNAs have also been shown to target RNA viruses (Skalsky and Cullen, 2010). The temporal variation in miRNA expression upon infection presented here will be informative in future studies on the interaction between OrV and miRNAs.

Thirdly, we observed a big downregulation of 26G RNAs at 38 hpi, which could be explained by competition between virus-derived and endogenous sRNAs, as competition between endogenous and exogenous sRNAs has been shown to affect the efficiency of the RNAi response, can force induction of one pathway over another, and can regulate gene expression (Kennedy et al., 2004; Sarkies et al., 2013; Yigit et al., 2006). Our results thus support these studies and suggest that activation of the DCR-1, DRH-1 and RDE-4 complex (Consalvo et al. 2024), which processes viral RNAs to form 23 nt RNAs, results in downregulation of the DCR-1, ERGO-1 and RRF-3 complex, which processes endogenous RNAs to form 26G RNAs (Vasconcelos Almeida et al. 2019). In agreement with this, it has been shown that the RNAi response to the virus redirects the Argonaute protein RDE-1 from its endogenous sRNA cofactors to antiviral function, leading to loss of repression of endogenous RDE-1 targets (Sarkies et al., 2013).

Altogether, these results contribute to our understanding of viral replication dynamics, miRNA expression upon infection, and the regulation of the RNAi pathway.

### Concluding remark

We report here a phenomenon taking place within infection of *C. elegans* with OrV, superinfection exclusion, whereby exposure to a primary infection limited viral replication during a secondary infection of the same strain or a closely related strain of OrV. Although our understanding of superinfection exclusion remains far from complete, the discovery of this phenomenon in a new pathosystem will enable future avenues of research.

## MATERIALS AND METHODS

### *C. elegans* strain maintenance and synchronization

Strains were maintained on nematode growth medium (NGM) plates seeded with *Escherichia coli* OP50 under standard conditions (Brenner, 1974; Stiernagle, 2006). N2 Bristol and the natural isolate JU2634 were maintained at 20 °C and SFE4 and SFE12, which are derived from N2, were maintained at 15 °C. *C. elegans* strains used in this study are listed in Suppl. Table S6.

To generate synchronized animal populations, plates containing eggs were meticulously rinsed with M9 buffer (0.22 M KH_2_PO_4_, 0.42 M Na_2_HPO_4_, 0.85 M NaCl, 0.001 M MgSO_4_) to eliminate larvae and adult animals while retaining the eggs. After 1 h, plates were subjected to a second M9 buffer wash to gather the larvae that had hatched during that period.

### OrV stock preparation, quantification and inoculation

JU2624 animals coming from one freshly starved plate were inoculated with OrV strains JUv1580 or JUv2572 by soaking in viral stocks kindly provided by Prof. M.A. Félix (Félix et al., 2011) for 1 h. The volume was then divided among 30 9 cm NGM plates and allowed to grow for 5 d until starved. Animals were then resuspended in 15 mL of M9, let stand for 15 min at room temperature, vortexed and centrifuged for 2 min at 400 g. The supernatant was centrifuged twice at 21,000 g for 5 min and then passed through a 0.2 μm filter.

RNA of the resulting viral stock was extracted using the Viral RNA Isolation kit (NYZ tech). The concentration of viral RNA was then determined by RT-qPCR (details below) and normalized using a standard curve. Primers used can be found in Suppl. Table S6.

For the standard curve cDNA of JUv1580 was obtained using Accuscript High Fidelity Reverse Transcriptase (Agilent) and reverse primers at the 3’ end of the RNA2 of the virus. Approximately 1000 bp of the 3’ end were amplified using a forward primer containing 20 bp coding the T7 promoter and DreamTaq DNA Polymerase (Thermo Fisher). The PCR product was gel purified using MSB Spin PCRapace (Invitek Molecular) and an *in vitro* transcription was performed using T7 Polymerase (Merck). The remaining DNA was then degraded using DNAse I (Life Technologies). RNA concentration was determined by NanoDrop (Thermo Fisher) and the number of molecules per µL was determined using the online tool EndMemo RNA Copy Number Calculator (https://www.endmemo.com/bio/dnacopynum.php). Primers used can be found in Suppl. Table S6.

Synchronized animal populations were inoculated with 2.8×10^9^ copies of JUv1580 or JUv2572 by pipetting the viral stock on top of the bacterial lawn containing the animals. Five hundred animals per plate were grown for RT-qPCRs, smiFISH, and for the transcriptomic analysis, except for the RNA-seq time-point of 12 hph, for which 2400 animals per plate were grown. Thirty animals per plate were used for survival assays.

### RNA extractions and RT-qPCRs

Inoculated and control animals were collected at the designated times with PBS 0.05% Tween. Samples were centrifuged for 2 min at 1350 rpm and the supernatant was discarded. Another two wash steps were performed before freezing the samples in liquid nitrogen. 500 µL of Trizol (Invitrogen) were added to the animal pellet and the pellet was disrupted by following five cycles of freeze-thawing and five cycles of 30 s of vortex followed by 30 s of rest. 100 µL of chloroform were then added and the tubes were shaken for 15 s and let rest for 2 min. Samples were centrifuged for 15 min at 11,000 g at 4 °C and the top layer containing the RNA was then mixed with the same volume of 100% ethanol. The sample was then loaded into RNA Clean & Concentrator columns (Zymo Research) and the rest of the protocol was followed according to manufacturer instructions.

RT-qPCRs were performed using Power SYBR Green PCR Master Mix (Applied Biosystems) on an ABI StepOne Plus Real-time PCR System (Applied Biosystems). 10 ng of total RNA were loaded and a standard curve (details above) was used for OrV quantifications. Primers used for RT-qPCRs can be found in Suppl. Table S6.

### smiFISH

The smiFISH protocol was adapted from previous studies (Parker et al., 2021; Tsanov et al., 2016). Inoculated and control animals were washed with PBS 0.05% Tween and centrifuged for 2 min at 1350 rpm four times discarding the supernatants. 800 µL of Bouins fixation mix (400 µL Bouins Fix, 400 µL methanol, 10 µL β-mercaptoethanol) were added and the sample incubated at room temperature for 30 min in a rotatory shaker. Samples were then frozen in liquid nitrogen and kept at −80 °C overnight. The following day, samples were gently shaken at 4 °C for 30 min and washed four times with borate Triton solution (20 mM H_3_BO_3_, 10 mM NaOH, 0.5% Triton) and five times with borate Triton 2% β-mercaptoethanol solution, leaving the borate Triton β-mercaptoethanol incubate for 1 h between each wash in the last three washes. The sample was then incubated for 5 min with wash buffer A (1 mL Stellaris RNA FISH Wash Buffer A, 1 mL deionized formamide and 3 mL H_2_O).

Forty probes against OrV were designed using Oligostan (Tsanov et al., 2016), 22 against RNA1 and 18 against RNA2. The sequences of the probes can be found in Suppl. Table S6. The probes were annealed as follows: 2 µL 0.83 µM probe set, 1 µL 50 µM FLAP-label, 1 µL NEB3 and 6 µL DEPC H_2_O were incubated at 85 °C for 3 min, 65 °C for 3 min and 25 °C for 5 min. The FLAP labels contained CAL Fluor 610 or Quasar 670 modifications at both the 5’ and 3’ ends of the following sequence: 5’ AATGCATGTCGACGAGGTCCGAGTGT (Biosearch Technologies).

One µL of annealed probe solution targeting RNA1 and 1 µL of annealed probe solution targeting RNA2 was then mixed with 98 µL of hybridization buffer (100 µL Stellaris FISH hybridization buffer, 25 µL Deionized formamide). 100 µL of hybridization buffer containing the FISH probes were then added to the samples and the samples were incubated overnight at 37 °C in a rotary shaker. The following day, samples were washed with wash buffer A2 (1 mL, Stellaris RNA FISH Wash Buffer A, 4 mL H_2_O), incubated for 30 min in wash buffer A2 at 37 °C, and then incubated for 30 min in Wash buffer A2 containing 25 ng of DAPI at 37 °C. Samples were then incubated 5 minutes in Stellaris RNA FISH Wash Buffer B, centrifuged and resuspended in a small volume of Stellaris RNA FISH Wash Buffer B. 0.1 ng of DAPI were added to each sample and samples were mounted using N-propyl gallate mounting medium. Samples were imaged using Leica Dmi8 microscope with Leica DFC9000 GTC sCMOS camera and objective HC PL APO 40×/0.95 CORR PH2. Images were analyzed and processed using ImageJ (FIJI) (Schindelin et al., 2012).

### Temporal dynamics of *pals-5p*::*GFP* fluorescence

Inoculated and control animals were placed in a 3% agarose pad containing 20 mM of sodium azide (NaN_3_) at the designated times and imaged using a Leica MZ10F fluorescence stereomicroscope with Leica Flexacam C3 camera, objective 10×/23B and mCherry M10F/MZ FLII and GFP3 MZ10F filters. Images were analyzed and processed using ImageJ (FIJI) (Schindelin et al., 2012). A ROI of the anterior intestine – from the start of the intestine to just before the vulva – was drawn for each worm and the average GFP intensity of the ROI was recorded. Fluorescence intensities of infected animals were transformed into *z*-scores using the mean and standard deviation of the intensity measured for control animals. Animals were classified as infected when the *z-*scores had a significant 1-tail *P* value.

### Survival assays

Synchronized inoculated and control animals were allowed to develop for 48 h at 25 °C. Adult worms were transferred to new plates every day until the end of egg laying, at which point they were allowed to stay on the same plate until the end of the assay. Animals were scored for survival every 24 h and scored as dead if they failed to respond when the nose and tail were touched three times and pharyngeal pumping had ceased.

### RNA preparation for RNA-seq

Library preparation and Illumina sequencing was done by Novogene Europe (https://www.novogene.com) using a NovaSeq 6000 platform and a Lnc-stranded mRNA-seq library method, ribosomal RNA depletion and directional library preparation, 150 paired end, and 6 Gb raw data per sample. Novogene checked the quality of the libraries using a Qubit 4 Fluorometer (Thermo Fisher Scientific), qPCR for quantification and Bioanalyzer for size distribution detection.

### RNA-seq data preprocessing

The quality of the raw paired-end data (fastq files) was assessed with FastQC (Andrews, 2010), which was summarised with MultiQC (Ewels et al., 2016). The cleaning of the reads was done with BBDuk The cleaning of the reads was done with BBDuk (Bushnell, 2024) (parameters *forcetrimleft = 10, ktrim = r, k = 31, mink = 11, qtrim = rl, trimq = 10, maq = 5, minlength = 60,* and), where adapter sequences were removed, the first ten 5’ nucleotides of each read were hard-clipped, nucleotides of poor quality (quality less than 10) were removed from the 3’ end, and reads less than 60 nucleotides were removed from the dataset. The library sizes varied from around 2M pairs to around 23M. However, C_12h_2 had almost 90 M, so a subset of this library was generated with Seqtk (parameters *sample -s100 19000000*) (https://github.com/lh3/seqtk).

### Differential expression analysis and functional enrichment analysis

The cleaned RNA-seq data was mapped to a reference genome consisting of *C. elegans* N2 genome assembly WBcel235 (GCF_000002985.6) and OrV strain JUv1580 vlc (Castiglioni et al., 2024a) using STAR version 2.7.11b (Dobin et al., 2013) and the read counts per gene were obtained within the same runs (parameters *--runMode alignReads --quantMode GeneCounts*).

DESeq2 version 1.40.2 (Love et al., 2014) was used to obtain the differentially expressed genes (DEG). We used two strategies: (*i*) the time series points were analysed as individual pairwise comparisons using Wald’s method and (*ii*) the epistatic effect of the super-infection was determined by following a mean reference model, as described in Law et al. (2020). The design tested by Wald’s method was superinfection *vs*. the sum of the single infections with the non-infected at 50 hph as the control. In all cases, a filter was applied to only include groups/genes with at least five reads per row in total. Furthermore, all log_2_-fold changes (*FC*) were shrunk using the *ashr* method (Stephens, 2017). DEGs were considered significant with a Benjamini and Hochberg false discovery rate (FDR) adjusted *P* < 0.05 and a log_2_*FC* ≥ 1.

### Viral load determined with RNA-seq

The mapped read information generated by STAR was used to calculate the viral load as the total number of reads mapped to the two viral chromosomes divided by their combined total length.

### Small RNA-seq analysis

Adapters were removed and sequences with indeterminacies were filtered using FASTP version 0.23.2 (Chen et al., 2018). Trimmed sequences between 15 and 40 nucleotides in length were aligned to the *C. elegans* genome (GCA_000002985.6) using Bowtie version 1.3.1 (Langmead et al., 2009) with parameters *--best -v 1 -k 1*. Sequences that aligned to the genome were quantified obtaining both absolute count and RPM matrices, which were used for differential expression analysis and visualization purposes, respectively. Reads with low counts (fewer than five counts in at least three samples) were removed from the absolute count matrix. Raw counts were then normalized with the median of ratios method and used for differential expression analysis using DESeq2 version 1.42.1 (Love et al., 2014). The Wald test was performed for hypothesis testing and the Benjamini and Hochberg method to control FDR. Sequences with an adjusted *P* < 0.05 were considered differentially expressed. The log_2_*FC* of sequences were shrunk using the *lfcShrink* function in DESeq2 to improve their accuracy.

Trimmed sequences of each library were annotated with Unitas version 1.8.0 (Gebert et al., 2017), with parameters *-species caenorhabditis_elegans -skip_dust off -trim_minlength 15 -trim_maxlength 40 -species_miR_only -na_for_phasi -tail 0*. Only the first annotation per sequence was considered for each sample, which were subsequently combined into a single file of unique sequences. The miRNA, rRNA, snRNA, and snoRNA annotation categories remained unchanged. However, the other annotation groups provided by Unitas were modified or reorganized. Sequences annotated as 5’ tRFs, 5’ tR-halves, 3’ tRFs, 3’ CCA-tRFs, 3’ tR-halves, tRF1, tRNA-leader, and misc-tRFs were grouped into tsRNA. Sequences of 23 nt that were not assigned to any of the previously mentioned categories were classified as 23 nt RNAs. Similarly, those sequences of 26 nt in length with a 5’ guanosine bias, whether unannotated or previously annotated as ncRNA, were classified as 26G RNAs. The remaining sequences were assigned to the category Other.

### miRNA-target interaction network

To construct a miRNA-target interaction network, pairs of miRNA and target genes were obtained from TarBase v9.0 (accessed on December 2024), a database of experimentally supported miRNA-target interactions. Only those pairs in which both the miRNA and the target gene were significantly differentially expressed (adjusted *P* < 0.05) with a |log_2_*FC*| > 0.5 were used. The up or downregulated status of all sequences aligning to each miRNA was considered to determine the status of said miRNA. To focus exclusively on canonical miRNA-mediated gene regulation, only miRNA-target pairs with opposite status were selected. The network was visualized using Cytoscape version 3.10.3.

### Other analysis and data representation

To determine the overlap between significant DEG groups, UpSet plots were performed with UpsetR v1.4.0 (Conway et al., 2017). Enrichment analyses of “Biological Process”, “Molecular Function”, “Cellular Component”, and KEGG pathways were conducted on significant DEGs using clusterProfiler v4.8.3 (Wu et al., 2021). Plots were created using ggplot2 v3.5.1 (Wickham, 2016). Other R libraries used for data processing and plot refinements were tidyr (Wickham et al., 2024), ggpattern (available at https://github.com/coolbutuseless/ggpattern) and ggbreak (Xu et al., 2021).

### Strain-specific OrV amplification

cDNA was made using total RNA extractions (details above) and Accuscript High Fidelity First Strand cDNA Synthesis Kit (Agilent). Two µL of cDNA were used for TaqMan SNP Genotyping (Thermo Fisher). The following sequences were used as reporter sequences: 5’ CAATTGCCAATCTC and 5’ CTTTGCCAGTGGTCCAG for RNA1 and RNA2 of JUv1580 and 5’ CTCCAATTGCCTATCTC and 5’ CTTTGCCAGTAGTCCAG for RNA1 and RNA2 of JUv2572. The reporter dyes VIC and FAM were used for JUv1580 and JUv2572, respectively, and NFQ was used as reporter quencher.

### Statistical analysis

Statistical analyses were performed using R version 4.4.2 in RStudio version 2024.09.1+394 (RStudio Team) or SPSS version 29.0.2.0 (IBM Corporation). For population comparisons, a Shapiro test of normality was first performed to determine if the data was sampled from a Gaussian distribution. Bartlett’s test was used to assess homogeneity of variance. For homoscedastic data drawn from a Gaussian distribution, comparisons were done using a one-way ANOVA. For data violating the ANOVA assumptions, non-parametric Kruskal-Wallis test was used. Pairwise Bonferroni’s *post hoc* tests were done. ANCOVA was used to compare viral load over time across multiple conditions, followed by *post hoc* Bonferroni’s tests for pairwise comparisons. Data from the survival assay was fitted to Kaplan-Meier regression model, followed up by a likelihood ratio test of significance.

Tests of significance used and sample sizes are indicated in the figure legends. No statistical method was used to pre-determine sample sizes. No samples or animals were excluded from analysis. The experiments were not randomized, and the investigators were not blinded to allocation during experiments and outcome assessment.

## ACKNOWLEDGMENTS

We thank Francisca de la Iglesia for excellent technical support and the members of the EvolSysVir lab for valuable comments and fruitful discussions. We also thank Wormbase (Davis et al., 2022). Some strains were provided by the Caenorhabditis Genetics Center, which is funded by NIH Office of Research Infrastructure Programs (P40 OD010440). Computations were performed on the HPC cluster Garnatxa at the Institute for Integrative Systems Biology (I2SysBio). This work was supported by grants PID2022-136912NB-I00 funded by MCIU/AEI/10.13039/501100011033 and by “ERDF a way of making Europe”, and CIPROM/2022/59 funded by Generalitat Valenciana to S.F.E. V.G.C. was supported by grant FJC2021-047264-I funded by MCIU/AEI/10.13039/501100011033 and by NextGenerationEU/PRTR.

## AUTHOR CONTRIBUTIONS

V.G.C: Writing - original draft, Conceptualization, Investigation, Writing - review & editing, Methodology, Resources, Funding acquisition, Validation, Supervision, Formal analysis, Project administration, Visualization. A.V-G: Investigation. A.G-S: Investigation, Methodology, Formal analysis, Visualization, Writing - review & editing. C.T: Investigation, Methodology, Formal analysis, Visualization, Writing - review & editing. G.G.G: Supervision, Writing - review & editing. S.F.E: Writing - original draft, Conceptualization, Writing - review & editing, Methodology, Resources, Funding acquisition, Data curation, Validation, Supervision, Formal analysis, Project administration.

## SUPPLEMENTARY INFORMATION

**Supplementary Figure S1.**
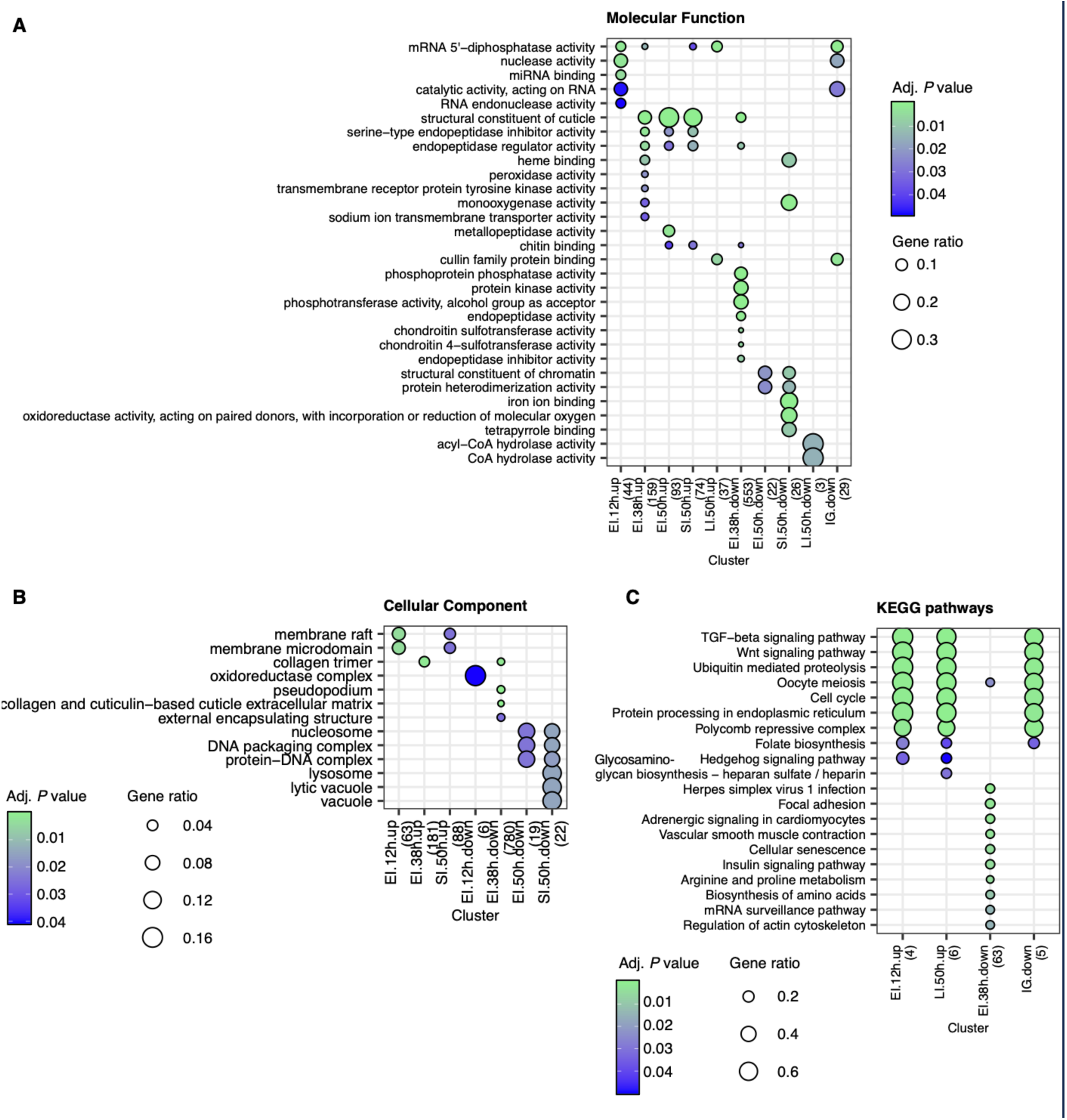
Functional enrichments upon superinfection. Details of the conditions in Fig. 3A. Only significant terms are shown (adjusted *P* ≤ 0.05). (A) GO:MF. (B) GO:CC. (C) KEGG pathways.

**Supplementary Figure S2.**
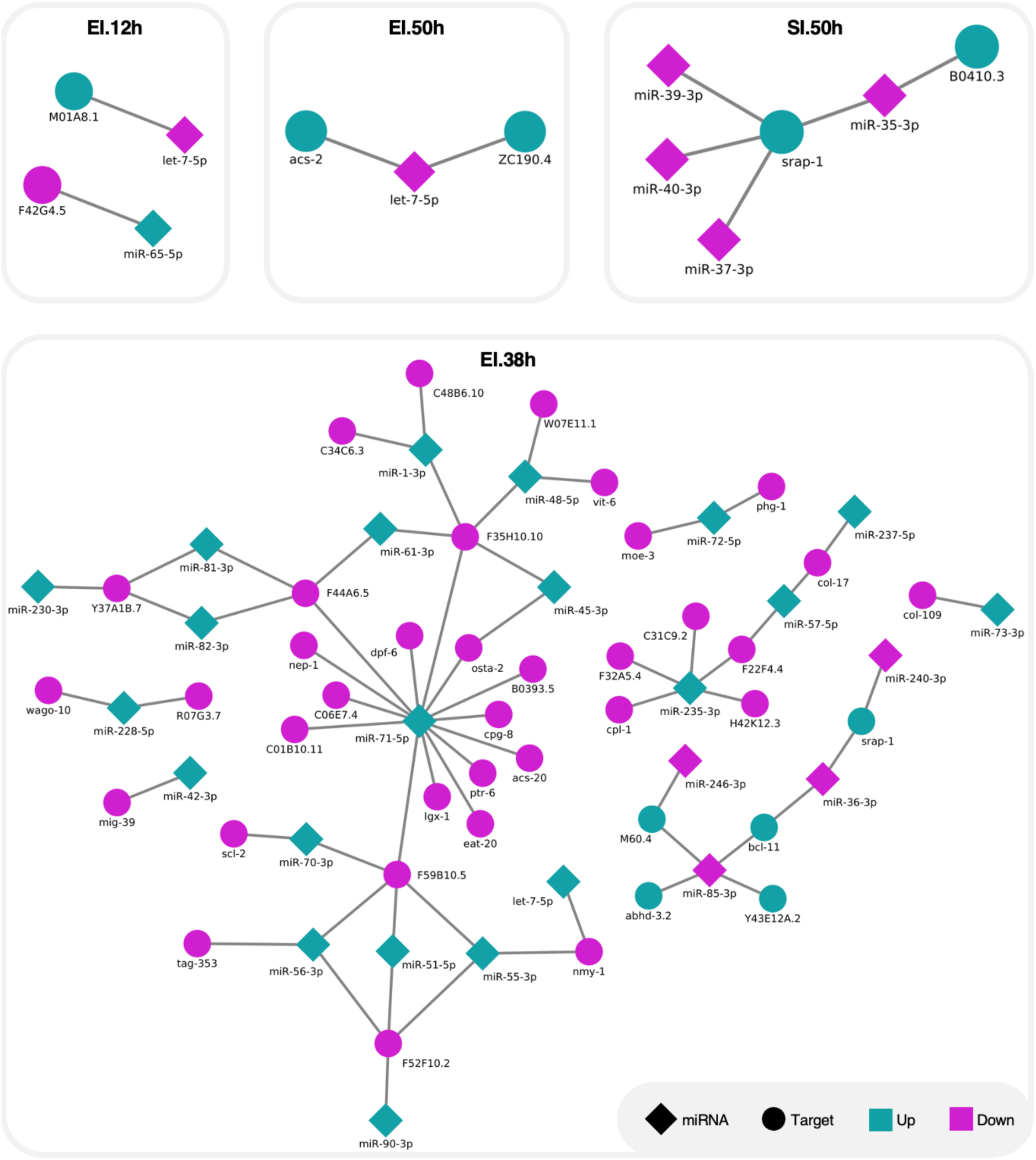
miRNA-target interaction network. Rhombuses and circles represent miRNAs and targets, respectively. Cyan and magenta represent up-regulated and down-regulated miRNA or targets.

**Supplementary Figure S3.**
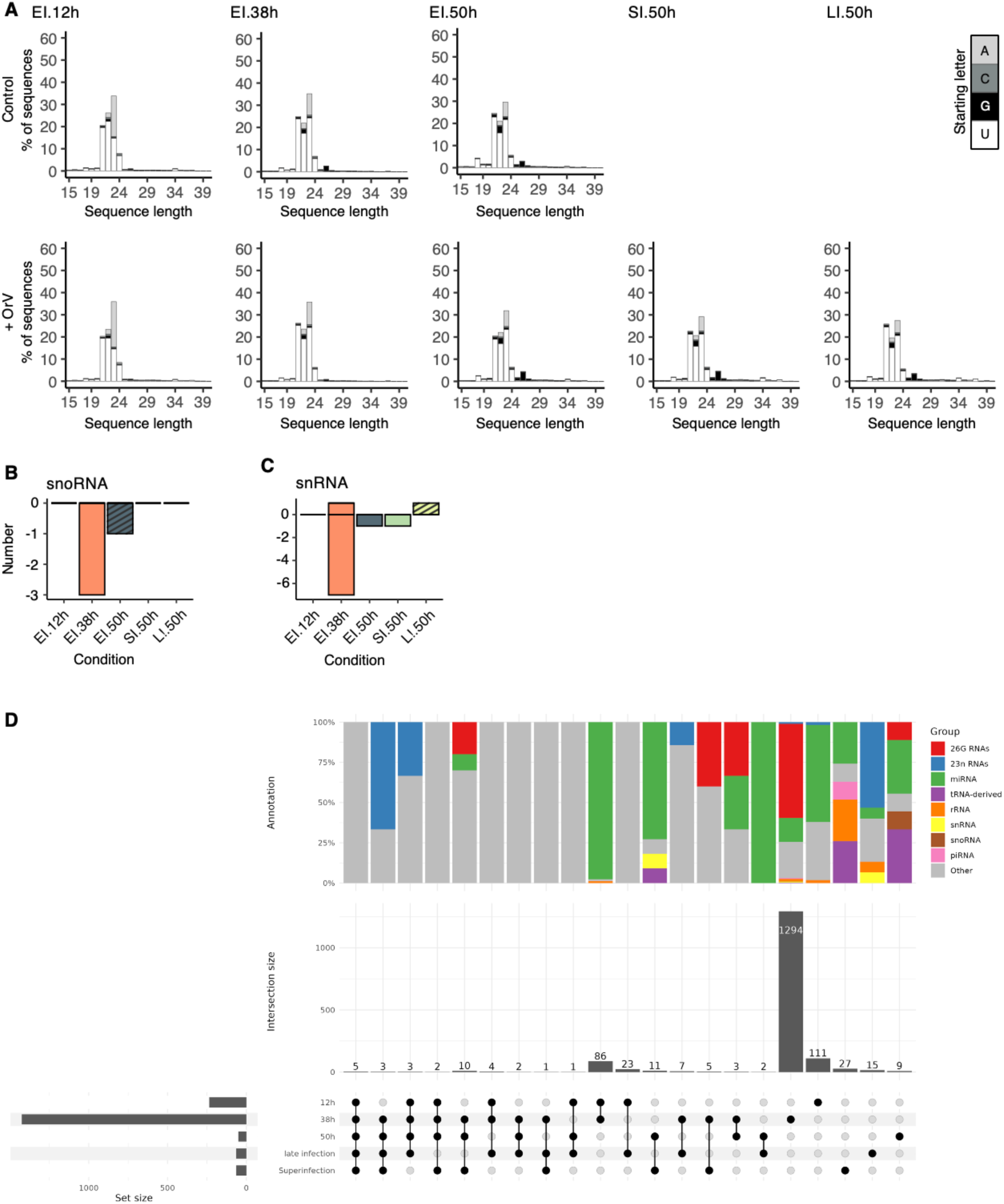
Functional enrichments upon superinfection. Details of the conditions in Fig. 4. (A) Percentage of fragments of each sequence length in each condition. (B) Number of upregulated and downregulated snoRNAs in each condition. (C) Number of upregulated and downregulated snRNAs in each condition. (D) Upset plot showing the intersections between the DEFs in each condition and the classes of small RNAs within each intersection. The points indicate the sets that are counted in the vertical bars. The horizontal bars indicate the total number of DEFs in each condition.

